# HIV-1 Vif disrupts phosphatase feedback regulation at the kinetochore, leading to a pronounced pseudo-metaphase arrest

**DOI:** 10.1101/2024.07.30.605839

**Authors:** Dhaval Ghone, Edward L. Evans, Madison Bandini, Kaelyn G. Stephenson, Nathan M. Sherer, Aussie Suzuki

**Author notes:** Corresponding authors: Aussie Suzuki and Nathan Sherer. These authors contributed equally (Dhaval Ghone, Edward L. Evans). Laboratory for Optical and Computational Instrumentation, University of Wisconsin-Madison, Madison, Wisconsin, 53705, USA.

## Abstract

Virion Infectivity Factor (Vif) of the Human Immunodeficiency Virus type 1 (HIV-1) targets and degrades cellular APOBEC3 proteins, key regulators of intrinsic and innate antiretroviral immune responses, thereby facilitating HIV-1 infection. While Vif’s role in degrading APOBEC3G is well-studied, Vif is also known to cause cell cycle arrest, but the detailed nature of Vif’s effects on the cell cycle has yet to be delineated. In this study, we employed high-temporal single-cell live imaging and super-resolution microscopy to monitor individual cells during Vif-induced cell cycle arrest. Our findings reveal that Vif does not affect the G2/M boundary as previously thought. Instead, Vif triggers a unique and robust pseudo-metaphase arrest, distinct from the mild prometaphase arrest induced by Vpr. During this arrest, chromosomes align properly and form the metaphase plate, but later lose alignment, resulting in polar chromosomes. Notably, Vif, unlike Vpr, significantly reduces the levels of both Protein Phosphatase 1 (PP1) and 2A (PP2A) at kinetochores, which regulate chromosome-microtubule interactions. These results unveil a novel role for Vif in kinetochore regulation that governs the spatial organization of chromosomes during mitosis.

## Introduction

The human immunodeficiency virus type 1 (HIV-1) weakens the immune system by depleting CD4+ T cells, eventually causing the Acquired Immunodeficiency Syndrome (AIDS) (Deeks et al, 2015; Swanstrom & Coffin, 2012). As a result, HIV-1 infected individuals have an increased susceptibility to specific cancers and other health complications (Cohen et al, 2016; Grulich et al, 2007; Hernández-Ramírez et al, 2017; Parkin, 2006). After HIV-1 enters a host cell, its RNA genome undergoes reverse transcription to form double-stranded DNA, followed by integration of the DNA provirus into the host’s genome. Using the host’s transcriptional machinery, HIV-1 transcribes its genome into spliced, partially spliced and completely unspliced viral mRNAs, facilitating viral gene expression and infectious virion production (Jouvenet et al, 2009; Nguyen & Hildreth, 2000; Ono & Freed, 2001). While the mechanisms underlying CD4+ T cell depletion during HIV-1 infection remain an active area of research, evidence suggests that both direct cytopathic effects of HIV-1 and chronic hyperactivation of the immune system contribute significantly. These processes drive apoptosis and induce pyroptosis in CD4+ T cells, leading to their progressive loss (Doitsh & Greene, 2016; Vidya Vijayan *et al*, 2017).

HIV-1 encodes four accessory viral proteins (Vif, Vpr, Vpu, and Nef) that are nonessential for virus replication in some ex vivo cell culture systems (Gabuzda et al, 1992) but play crucial immunomodulatory roles in vivo (Malim & Emerman, 2008). The primary role of Vif (Virion Infectivity Factor) is to facilitate the proteasomal degradation of APOBEC3 (A3) family of cytidine deaminases (e.g., A3F, A3G, and A3H). A3 proteins introduce deleterious mutations into the HIV-1 genome by deaminating cytosine residues in the viral single-stranded DNA during reverse-transcription, converting them to uracil (Chiu & Greene, 2009; Okada & Iwatani, 2016). Vif orchestrates A3 protein degradation by recruiting an E3 ubiquitin ligase complex (Conticello et al, 2003; Marin et al, 2003; Sheehy et al, 2003; Stopak et al, 2003; Yu et al, 2003). This degradation prevents A3 proteins from being incorporated into budding viral particles, ensuring that the progeny virions remain infectious.

Independently of its primary role of A3 protein degradation (DeHart et al, 2008; Du et al, 2019; Zhao et al, 2015), several studies have shown Vif to induce cell cycle arrest and cell death in CD4+ T cells and several other cell types (DeHart et al., 2008; Du et al., 2019; Greenwood et al, 2016; Marelli et al, 2020; Nagata et al, 2020; Sakai et al, 2006; Salamango et al, 2019; Zhao et al., 2015). However, the molecular mechanisms that underpin these effects remain unclear. A prior study indicated that p53, a tumor suppressor, is required for Vif-induced G2/M cell cycle arrest (Izumi et al, 2010). Several recent studies suggest that Vif’s cell cycle arrest activity is due to the loss of B56 proteins that are regulatory subunits of protein phosphatase 2A (PP2A) (DeHart et al., 2008; Du et al., 2019; Greenwood et al., 2016; Marelli et al., 2020; Nagata et al., 2020; Salamango et al., 2019; Salamango et al, 2020; Zhao et al., 2015). The PP2A-B56 complex is a serine-threonine protein phosphatase that participates in various key G2 and mitotic processes (Lee et al, 2017; Schuhmacher et al, 2019). Additionally, earlier research has demonstrated that Vif can lead to reductions in Cyclin F (Augustine et al, 2017), a non-canonical cyclin critical for late S- and G2-phase progression (Clijsters et al, 2019; Enrico et al, 2021), or affect both Cdk1 and Cyclin B1, which are essential for the transition into and out of mitosis (Sakai et al, 2011).

These prior studies have predominantly employed flow cytometry-based techniques to measure cell cycle phase population densities. However, flow cytometry has limitations in its ability to differentiate between late S, G2, and M phases, because it categorizes cell cycle phases solely based on relative DNA content. Accordingly, in this study we prioritized high-temporal single-cell live imaging that would allow us to directly observe the disruptions of cell cycle triggered by Vif expression. We demonstrate that Vif induces a highly unique and robust pseudo-metaphase arrest, irrespective of cell line tested or p53 status. Additionally, we found that Vpr unexpectedly induces a distinct mitotic delay, clearly different from the pseudo-metaphase arrest caused by Vif. Vif, but not Vpr, disrupts the localizations of PP2A-B56 at the kinetochores during prometaphase, leading to a slight yet significant delay in the alignment of chromosome at metaphase. This disruption results in reduced localization of the Astrin-SKAP-PP1 complex at the kinetochores, causing improper kinetochore-microtubule binding affinity due to increased phosphorylation of the microtubule binding protein at kinetochores. These effects yield unbalanced forces between sister chromatids, resulting in misaligned chromosomes and abnormal chromosomal movements in association with spindle microtubules. These insights provide a deeper understanding of Vif’s impact on the regulation of the host cell cycle, a conserved feature of Vif that may have potential relevance to HIV-1 pathogenesis in vivo.

## Results

### Vif and Vpr induce distinct forms of mitotic arrest

Previous research demonstrated that both Vif and Vpr expression causes cell cycle arrest and cytotoxicity in CD4+ T cells and as well as many cancer cell lines (Augustine *et al*., 2017; Emerman, 1996; Evans *et al*, 2018; Nagata *et al*., 2020; Sakai *et al*., 2011; Sakai *et al*., 2006; Salamango *et al*., 2019; Salamango *et al*., 2020; Wang *et al*, 2011; Wang *et al*, 2008). To investigate the nature of the cell cycle arrest induced by Vif, we employed high-temporal live cell imaging using the triple negative breast cancer Cal51 cell line. This cell line was chosen for several reasons; it has been engineered for precise cell cycle tracking through CRISPR-Cas9-mediated endogenous tagging of Histone H2B with mScarlet, allowing visualization of DNA, and Tubulin with mNeonGreen, to enable monitoring the microtubule cytoskeleton (Scribano et al, 2021). This approach minimizes the confounding effects of exogenous overexpression of fluorescently labeled Histone and Tubulin proteins on cell cycle progression (**Fig 1A**). Moreover, Cal51 cells are well-suited for long-term live cell imaging assays due to their adherent growth properties, which facilitate extended single-cell monitoring. They also exhibit a stable, near-diploid karyotype and retain wild-type p53 expression, making them an ideal model for cell cycle research (Lynch et al, 2022). For these experiments, we infected cells with the NL4-3 strain HIV-1 reporter viruses expressing either Vif (“Vif”), Vpr (“Vpr”), a combination of both (“Vif+Vpr”), or a lack of both (“Control”) (**Fig S1A**) (Evans *et al*., 2018). Note that the NL4-3 strain encodes all known HIV-1 proteins and serves as a well-established model for studying various aspects of HIV-1 biology (Mustafa & Robinson, 1993). These reporter viruses contain cyan fluorescent protein (CFP), allowing us to identify infected cells under fluorescence microscopy. To focus our study on the effects of these proteins on the cell cycle, the reporter viruses do not express the viral Env and Nef proteins, which are known to exhibit cytotoxicity (Elder, 2002; Emerman, 1996; Evans *et al*., 2018).

**Figure 1:**
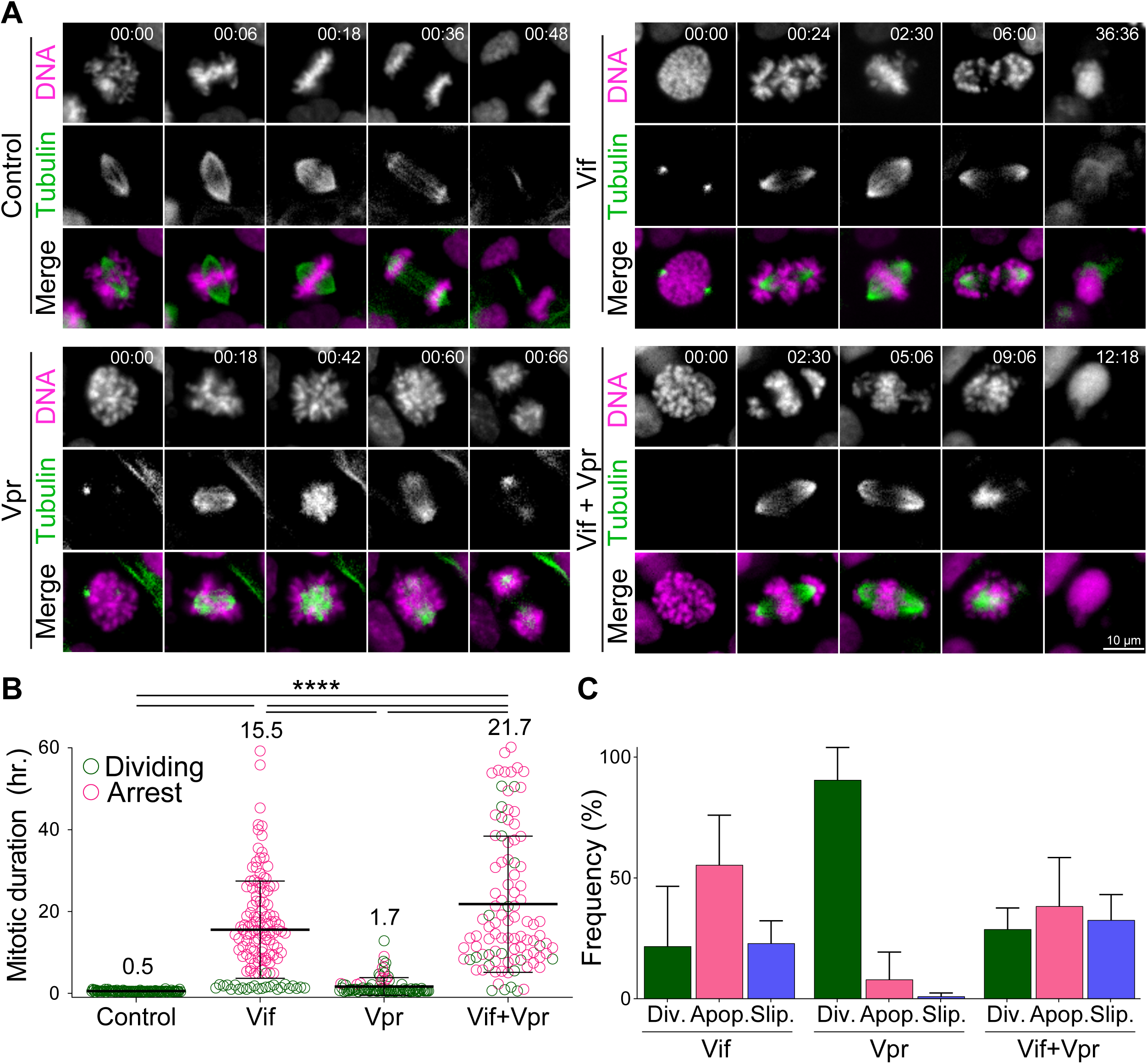
Vif and Vpr induce distinct forms of mitotic arrest. (A) Representative live cell image for Cal51 cells with H2B-mScarlet and Tubulin-mNeonGreen expressing Control, Vif, Vpr, or Vif+Vpr reporter viruses. (B) Average mitotic duration of Cal51 cells expressing respective reporter virus (n=100 for each from two replicates). (C) Frequency of cell fate after mitosis for Cal51 cells expressing respective reporter virus. (n=100 for each from two replicates)

We then assessed the mitotic duration defined as the time between nuclear envelope breakdown (NEBD) and anaphase onset in CFP-positive cells using video microscopy. Vif-expressing Cal51 cells demonstrated a prolonged mitosis of ∼16 hours, in contrast to only 30 min in Control cells (**Fig 1B and Movies S1-2**). Great majority of Vif-expressing cells eventually succumbed to apoptotic cell death or exhibited mitotic slippage, wherein the cell exited mitosis without completing chromosome segregation (**Fig 1C and S1B**).

Vpr has been known to induce G2/M arrest in a variety of cell types, as evidenced by flowcytometry (Bartz *et al*, 1996; Elder, 2002; Emerman, 1996; Hall *et al*, 2024; He *et al*, 1995; Jowett *et al*, 1995; Sakai *et al*., 2006). We next examined the differences in G2/M arrests induced by Vif and Vpr. While we expected Vpr to induce G2 phase arrest due to its abilities to cause DNA damage, our live-cell imaging revealed that Vpr-expressing cells also experienced a prolonged mitosis of ∼1.7 hours, which is significantly shorter than in Vif-expressing cells (**Fig 1A-B**). Interestingly, Vif+Vpr-expressing cells exhibited a prolonged mitosis lasting ∼21.7 hours, indicating that Vif plays a dominant role in mitotic arrest when both proteins are present. Supporting this, the majority of Vif-expressing or Vif+Vpr-expressing mitotic cells underwent apoptotic cell death, whereas Vpr-expressing mitotic cells either completed division or experienced mitotic slippage. In summary, although both Vif and Vpr can induce a prolonged mitosis, Vif causes significantly more severe mitotic arrest, leading to cell death.

To pinpoint the specific sub-stage of mitosis affected by Vif expression, we closely assessed chromosome alignment during metaphase. Notably, most Vif-expressing Cal51 cells successfully achieved metaphase chromosome alignment (metaphase plate) similar to the control cells, but with a slight delay, reaching it approximately 1.5 hours post-NEBD compared to the control’s ∼30 minutes. (**Fig 2A-C**). However, this alignment was unstable and deteriorated over time in Vif-expressing cells. Mitotic arrest induced by common mitotic inhibitors is seen in prometaphase before achieving the metaphase plate (Choi et al, 2011). However, the mitotic arrest caused by Vif was distinctive because cells were able to complete prometaphase but then gradually lost proper chromosome spatial organization at the metaphase plate over time. Accordingly, we termed this block “pseudo-metaphase arrest”. Consistent with our findings in Cal51 cells, other commonly used cell lines for cell cycle studies, such as HeLa and MDA-MB-231, also demonstrated significant mitotic arrest (approximately 12 hours for both) following Vif expression, which subsequently led to either apoptotic cell death or mitotic slippage (**Fig S1C-E and S2A-C**). Similar to Cal51, the majority of these Vif-expressing cells were able to establish a metaphase plate early but were unable to enter anaphase (**Fig S1F-G and S2D-E**). Consistent with these results, Vif-expressing HeLa cells exhibited a markedly higher mitotic index compared to Control cells at 72 hours post-infection in fixed immunofluorescence (IF) (**Fig S1H**). In conclusion, Vif triggers a marked pseudo-metaphase arrest in a range of cell lines. Most of these arrested cells experienced either apoptotic cell death or mitotic slippage, suggesting a conserved underlying mechanism.

**Figure 2:**
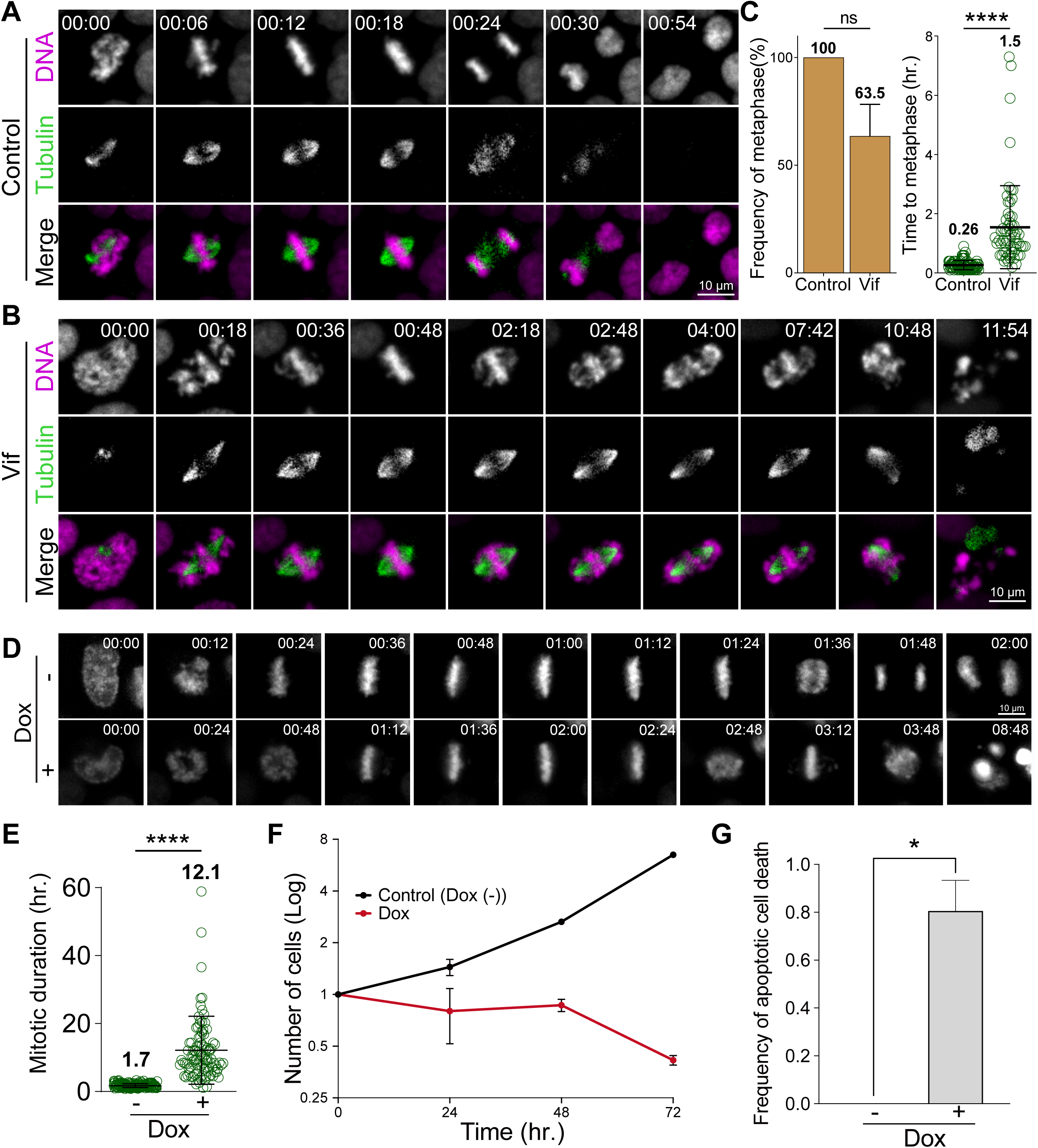
Vif induces robust pseudo-metaphase arrest. (A) Representative live cell images for Cal51 cells with H2B-mScarlet and Tubulin-mNeonGreen expressing Control reporter virus. (B) Representative live cell image for Cal51 cells expressing Vif reporter virus. (C) Frequency of cells that achieve metaphase plate and time taken to achieve metaphase plate for cells in (A) and (B) (n=100 for each from two replicates). (D) Representative live-cell images of Vif conditional expressed HeLa cell with or without Doxycycline. (E) Average mitotic duration in condition (D) (n=100 cells for each from two replicates). (F) Quantification of viable cells over time after Dox induction. (G) Quantification of apoptotic cells after Dox induction.

### Solo Vif expression is sufficient to trigger a robust pseudo-metaphase arrest

To determine if Vif expression alone is sufficient to induce a robust pseudo-metaphase arrest in the absence of other viral factors, we next engineered HeLa cells to conditionally express codon optimized Vif (CO-Vif) under the control of a doxycycline-inducible promoter (Das et al, 2016). As a control, we employed the same system but with mNeonGreen expression instead of Vif. Control cells displayed mNeonGreen signals approximately 10 hours post-doxycycline induction. In line with these expression kinetics, cells expressing CO-Vif almost invariably exhibited pseudo-metaphase arrest roughly 10 hours post-induction; with cells arrested for ∼15h in contrast to Control cells that proceeded through mitosis in ∼1 hour (**Fig 2D-E and Movies S3-4**). While Control cells continued to propagate, those expressing Vif did not, with Vif expression alone confirmed as sufficient to trigger prolonged pseudo-metaphase arrest and apoptotic cell death (**Fig 2F-G**).

### Vif accelerates G2 progression with no effect on the G1 or S phases

We next asked if Vif altered other stages of the cell cycle in addition to mitosis. To this end, we developed a novel method that allowed us to accurately distinguish between G1, S, and G2 phases in individual Cal51 reporter cells during live cell imaging based on tracking changes to the intensity of Histone H2B-mScarlet over time (see **Methods**). This method offers the advantage of allowing us to measure temporal changes of the DNA content at single cell resolution with high accuracy. Briefly, during S phase, H2B-mScarlet signals increased steadily, eventually plateauing and remaining constant throughout the G2 phase. **Fig 3A** presents example images and an intensity profile covering the period from the end of one mitosis to the beginning of the next in a Control Cal51 cell. Using this method, we observed no significant differences in the durations of either G1 or S phases in both Control or Vif-expressing cells. However, Vif-expressing cells exhibited a slight yet statistically significant reduction in G2 phase duration compared to Control cells (**Fig 3B-C and S3A**). Consistent to Cal51 cells, Vif expression also did not significantly impact the duration of interphase in two additional cell lines, RPE1 and MDA-MB-231 cells (**Fig. S3B).** In summary, these findings demonstrated that Vif expression induces pseudo-metaphase arrest without notably affecting the overall duration of interphase (the cumulative time of G1, S, and G2 phases).

**Figure 3:**
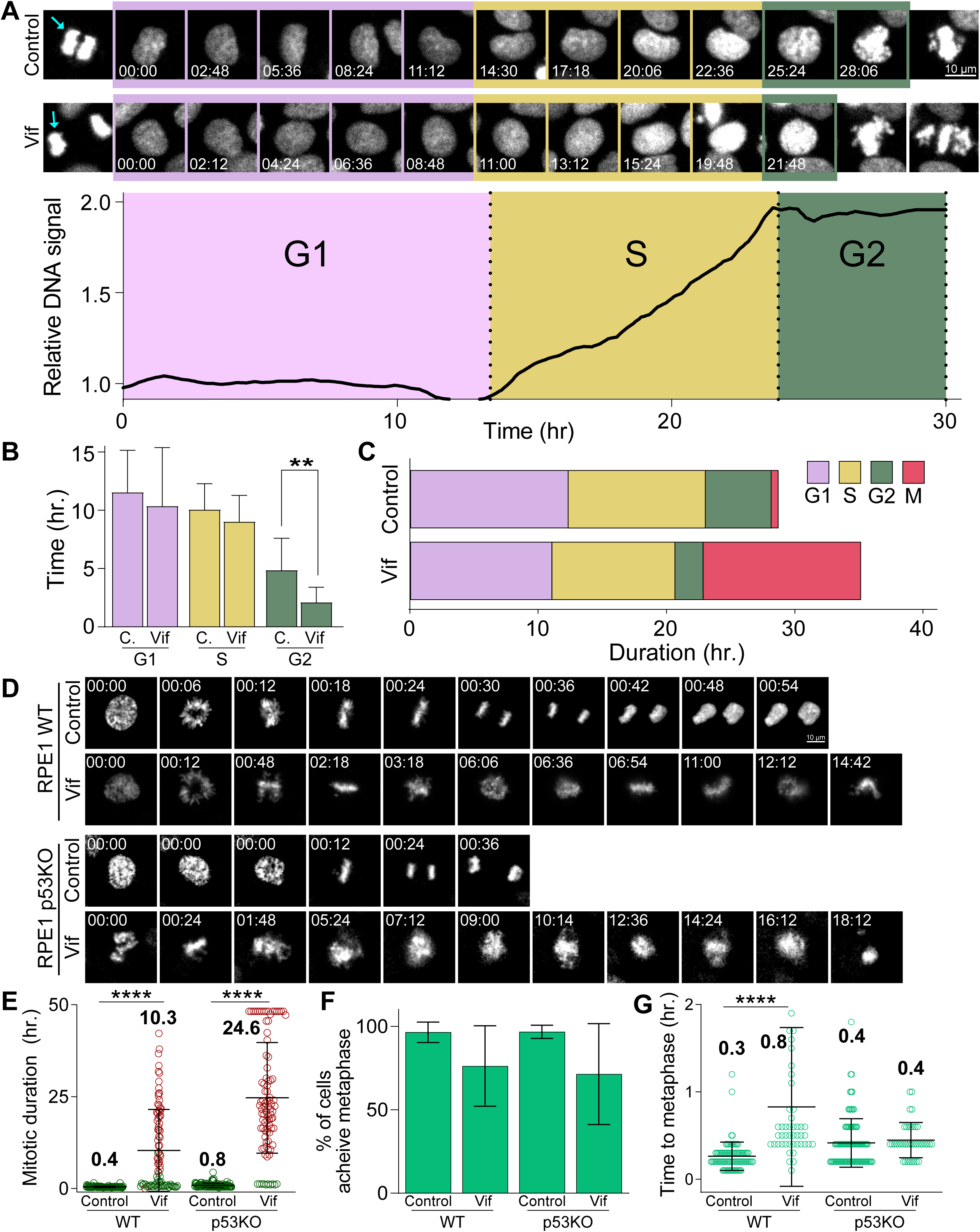
Vif does not alter G1 or S phase progression, accelerates G2 progression, and induces pseudo-metaphase arrest independent of p53. (A) Top: Representative image of Cal51 cells progressing through G1, S, and G2. Bottom: Representative trace for relative signal intensity of the nucleus through cell cycle. (B) Average duration of G1, S, and G2 phases in Control and Vif-expressing cells (n= 9 for Control and 11 for Vif, from two replicates). (C) Total cell cycle duration for Control and Vif-expressing cells. (D) Representative live cell images for Control and Vif-expressing WT or p35KO RPE1 cells. (E) Average mitotic duration in WT or p53KO RPE1 cells (n= >85 cells, from two replicates). (F) Frequency of cells which achieve metaphase plate for cells in (E). (G) Average time taken to achieve metaphase plate for cells in (F).

### Vif Induces Pseudo-Metaphase Arrest Independently of p53

A previous study indicated that Vif-induced cell cycle arrest is due to tumor suppressor p53 (Izumi et al., 2010), which is well established to trigger G2 cell cycle arrest in response to DNA damage (St Clair et al, 2004; Stark & Taylor, 2006; Taylor & Stark, 2001). Considering that we had already observed Vif inducing pseudo-metaphase arrest in cell lines with functionally inactivated p53, such as HeLa (Evans et al., 2018; Wang et al., 2011) and MDA-MB-231 (Olivier et al, 2002) (**Fig S1C-D and S2A-B**), we further investigated the potential p53-dependency by assessing Vif’s effects in p53 null knockout (p53 KO) RPE1 (Mardin et al, 2015) or HCT116 cell lines (**Fig 3D-G**). Both wild-type (p53 +/+) and p53 KO RPE1 and HCT116 cells demonstrated significant pseudo-metaphase arrest in response to viral Vif expression. Specifically, RPE1 wild-type cells were arrested for >10 hours, RPE1 p53 KO cells for ∼25 hours, and both HCT116 wild-type and p53 KO cells for >6 hours. In contrast, cells infected with the Vif-negative Control virus showed no delay in mitosis (∼30 min for both cell lines) (**Fig 3D-E and S3D-E**). All cell lines, regardless of p53 status, managed to establish a chromosome metaphase plate in the presence of Vif expression (**Fig 3F-G and S3F-G**). However, most Vif-expressing cells exhibited apoptotic cell death or mitotic slippage (**Fig S3C and S3H**).

### Vif-induced pseudo-metaphase arrest disrupts spatial organization of chromosomes and spindle poles

To further characterize the mitotic defects caused by Vif expression, we carefully assessed Vif’s effects on chromosome alignment at the metaphase plate. To this end, we employed super-resolution microscopy and stained for CENP-C, microtubules, and DNA (see **Methods**). CENP-C is a marker for kinetochores, the platform for microtubule attachment on mitotic chromosomes. Our findings revealed that ∼100% of Vif-expressing mitotic cells exhibited misaligned chromosomes, with the great majority of these misaligned chromosomes concentrated at spindle poles as polar chromosomes (**Fig 4A-B, S4A, and Movies S5-6**).

**Figure 4:**
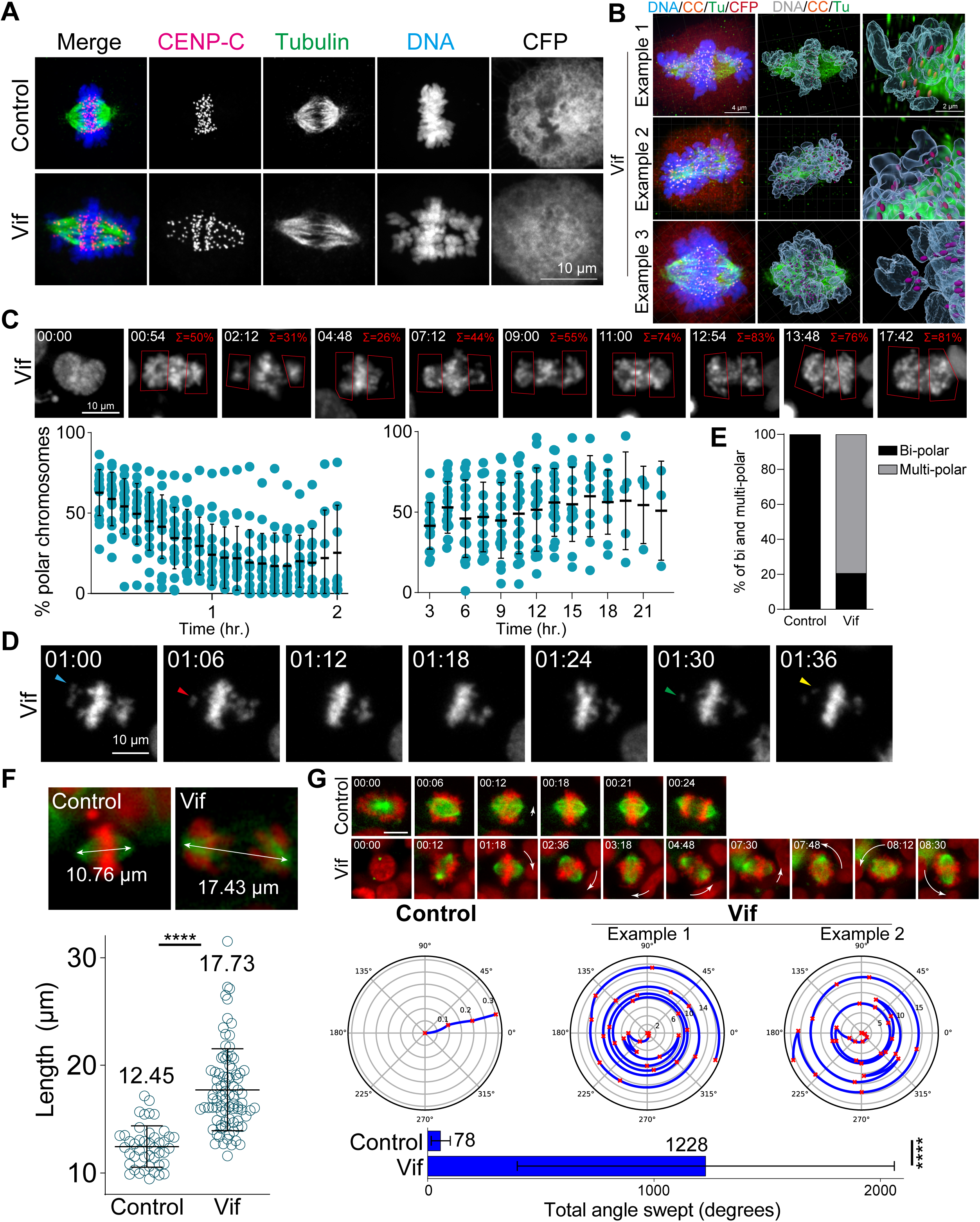
Vif induces polar chromosomes, multi-polar spindles, and abnormal chromosome movements. (A) Representative immunofluorescence images labeled for CENP-C (as a kinetochore marker), microtubule, and DNA in Control and Vif-expressing HeLa cells. (B) Example super-resolution images labeled for CENP-C, microtubule, and DNA in Vif-expressed HeLa cells showing polar chromosomes. (C) Representative live cell image of Vif-expressing cells where polar chromosomes were quantified by compartmentalizing polar regions. Bottom: Quantification of polar chromosome frequency overtime. (D) Representative high-temporal live cell images (6 minutes interval) showing rapid chromosome moving towards and away from the spindle poles. (E) Fraction of Cal51 cells showing abnormal number of poles at some point during mitosis. (F) Top: Representative images of maximum mitotic spindle length for Control and Vif-expressing Cal51 cells. Bottom: Average maximum mitotic spindle length of Control and Vif-expressing cells. (G) Top: Representative live cell image of Control and Vif-expressing Cal51 cells over time showing dynamic spindle spinning. Center: Representative figures showing relative orientation (angle) of the spindle axis over time (radius). Bottom: Average total angle swept during mitosis.

We next explored the dynamics of chromosome spatial organization in Vif-expressing cells by using live cell imaging. To do this, we first quantified the proportion of cells exhibiting polar chromosomes at any time point during metaphase/pseudo-metaphase for all four of our cell systems (HeLa, RPE1, MDA-MB-231, and Cal51 cells). Consistent with our fixed-cell analysis, we observed ∼100% of Vif-expressing cells exhibiting misaligned polar chromosomes at some time point during the prolonged mitosis, in contrast to Control cells in which misaligned chromosomes were only rarely observed (**Fig S4B**). To define the dynamic nature of chromosome movements, we segmented cells into two compartments, polar and equatorial, and then measured the Histone H2B-mScarlet signals within each of these compartments in Cal51 cells over time (**Fig 4C**). Vif-expressing cells exhibited an initial decrease in the frequency of polar chromosomes shortly after NEBD, but this frequency increased significantly during the extended pseudo-metaphase; with pronounced polar chromosomes comprising ∼50% of the total DNA. Notably, these misaligned chromosomes continuously oscillated between the poles and the metaphase plate, as shown in **Fig 4D**.

Consistent with abnormal chromosome dynamics, ∼25% of HeLa cells expressing Vif exhibited multi-polar spindles (>2 poles) at 72h post-infection based on fixed cell analysis (**Fig S4C**). To corroborate these findings, we used high-temporal live cell imaging to track and quantify spindle poles using mNeonGreen-Tubulin in Cal51 cells. We observed that ∼80% of cells expressing Vif demonstrated multi-polarity at some time point during the extended mitosis. Moreover, the number of spindle poles varied dramatically in arrested cells, ranging from a monopole to as many as five poles (**Fig 4E and S4D**).

The integrity of spindle poles is crucial for maintaining the position of the metaphase plate during mitotic progression, so that the length of microtubules making up the mitotic spindle is tightly regulated and typically remains stable until anaphase onset. Interestingly, we found that mitotic spindles in Vif-expressing cells were significantly stretched (∼18 µm in length) as compared to Control cells (∼12 µm) (**Fig 4F and S4E**). Moreover, although mitotic spindles are typically stationary, we observed spindles in Vif-expressing cells to exhibit dynamic spinning. To define these observations quantitatively, we measured the average angle swept by individual mitotic spindles over time in the presence or absence of Vif expression, observing a greater than 15-fold increase in the angle covered by spindles in Vif-expressing cells as compared to Control cells (**Fig 4G and S4F**). In summary, Vif induces dynamic movements in both chromosomes and spindle poles during extended pseudo-metaphase, resulting in severely misaligned polar chromosomes.

### Vif, but not Vpr, disrupts the proper localization of PP2A-B56 at the kinetochores

Microtubule assembly at the kinetochore is regulated by an intricate network of kinase and phosphatases (Saurin, 2018). PP2A-B56 is recruited to kinetochores during prometaphase, where it plays a crucial role in microtubule assembly and the proper alignment of chromosomes (Foley & Kapoor, 2013; Foley *et al*, 2011). Previous studies demonstrated that Vif can significantly degrade B56 proteins, as shown by western blots (Greenwood *et al*., 2016; Marelli *et al*., 2020; Nagata *et al*., 2020). Therefore, we investigated whether Vif-expressing mitotic cells had diminished B56 at kinetochores. To address this, we performed quantitative immunofluorescence (qIF) using specific antibodies against B56 and CENP-C (as a kinetochore marker) in Control, Vif-expressing, and Vpr-expressing cells. We found that B56 signals at kinetochores, regardless of aligned (equatorial) or unaligned (polar) chromosomes, were significantly reduced in Vif-expressing cells compared to Control and Vpr-expressing cells (**Fig 5A-C**). To determine whether Vif-expressing cells remained free of additional, non-kinetochore-bound pools of B56, we performed qIF in nocodazole treated cells. It has been demonstrated that nocodazole, microtubule depolymerizer, can enhance B56 kinetochore localization (Foley *et al*., 2011). As expected, Control cells showed further recruitment of B56 to kinetochores upon nocodazole treatment, whereas Vif-expressing cells did not (**Fig 5A and 5C**). These results suggest that Vif-mediated degradation of B56 is sufficient to significantly reduce B56 levels at kinetochores during prometaphase, while Vpr has no effect on B56 levels at kinetochores. To further validate these results, we performed qIF on Polo-like Kinase 1 (Plk1). Plk1 is a key cell cycle regulator with critical roles at kinetochores in proper mitotic progression (Colicino & Hehnly, 2018). It is known that Plk1 levels at kinetochores are regulated by PP2A-B56, and depletion of B56 causes increased levels of Plk1 at kinetochores, leading to improper microtubule attachments (Foley *et al*., 2011). As expected, Plk1 levels at kinetochores were significantly dropped in metaphase as compared to prometaphase in Control cells (**Fig 5D-E**). In Vif-expressing cells, while Plk1 levels at kinetochores in equatorial chromosomes were lower than those on polar chromosomes, Vif-expressing cells showed a global increase in Plk1 levels at kinetochores. More specifically, Plk1 levels at polar chromosomes in Vif-expressing cells were significantly higher than in Control prometaphase, and levels at aligned equatorial chromosomes were also significantly higher than in Control metaphase. In summary, Vif, but not Vpr, significantly diminishes PP2A-B56 levels at kinetochores, resulting in a delay of chromosome alignments.

**Figure 5:**
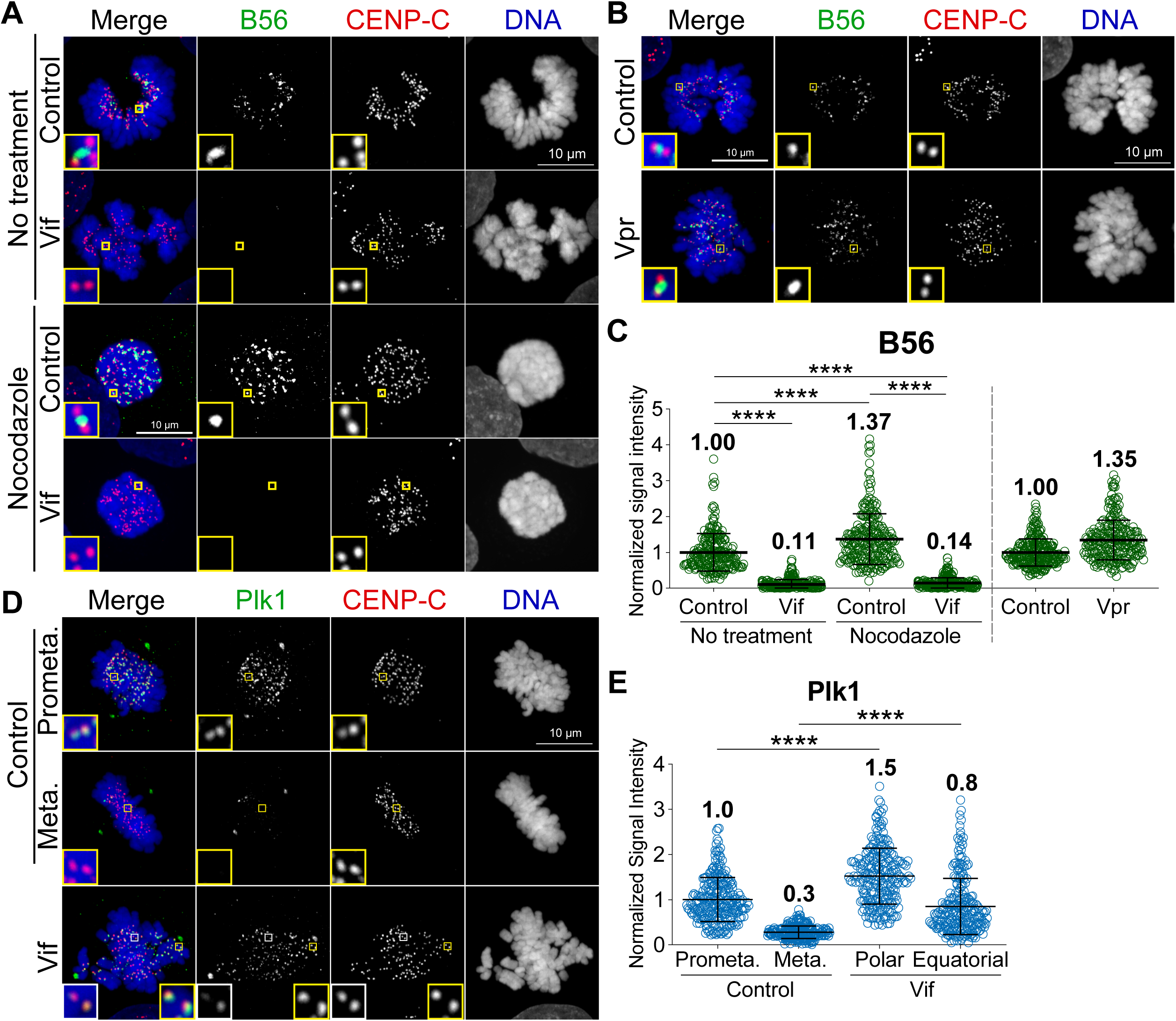
Vif, but not Vpr, disrupts the proper localization of PP2A-B56 at the kinetochores. (A) Representative maximum intensity projection of fixed Control and Vif-expressing HeLa cells with or without nocodazole treatment, immunofluorescently labeled for B56, CENP-C, and DNA. (B) Representative maximum intensity projection of fixed Control and Vpr-expressing HeLa cells, immunofluorescently labeled for B56, CENP-C, and DNA. (C) Average B56 level at kinetochores for cells in (A) and (B) (n=200 kinetochores over 8 cells from two independent replicates for each). (D) Representative fixed-cell immunofluorescent image of HeLa cells expressing Vif, immunofluorescently labeled for Plk1, CENP-C, and DNA. (E) Average Plk1 level at kinetochores for cells in (D) (n=200 kinetochores over 8 cells from two independent replicates for each).

### Vif impairs stable and balanced kinetochore microtubule attachments

We demonstrated that Vif-expressing cells exhibited abnormal dynamic chromosome movements (**Fig 4D-G**). Kinetochore-microtubule bindings are stabilized by both PP2-B56 and PP1 at kinetochores through an interplay and feedback mechanism (Saurin, 2018; Vallardi *et al*, 2017). Consequently, we hypothesized that the reduction of PP2A-B56 by Vif impaired the regulation PP1 phosphatase activities at kinetochores. To test this hypothesis, we quantified the levels of the Astrin-SKAP complex (hereinafter referred to as ‘Astrin’) at kinetochores by qIF in HeLa cells in the presence or absence of Vif expression. Astrin stabilizes kinetochore-microtubule attachments by recruiting PP1, which dephosphorylates Hec1, a microtubule binding protein at kinetochores, thereby promoting Hec1 binding to microtubules (Cheeseman *et al*, 2006; Conti *et al*, 2019; Dunsch *et al*, 2011; Manning *et al*, 2010; Schmidt *et al*, 2010; Zhang *et al*, 2015). As expected, Astrin signals at kinetochores significantly increased at metaphase compared to prometaphase in Control cells (**Fig 6A-B**). In contrast, Astrin levels at kinetochores on aligned chromosomes (equatorial) in Vif-expressing cells were approximately 50% of control, and Astrin levels on polar chromosomes largely undetectable (**Fig 6A-B**). We confirmed that levels of CENP-C, which is a core-structural kinetochore protein, did not change between Control and Vif-expressing cells, indicating that the reduction of Astrin in Vif-expressing cells was not due to compromised kinetochore integrity (**Fig 6A-B**).

**Figure 6:**
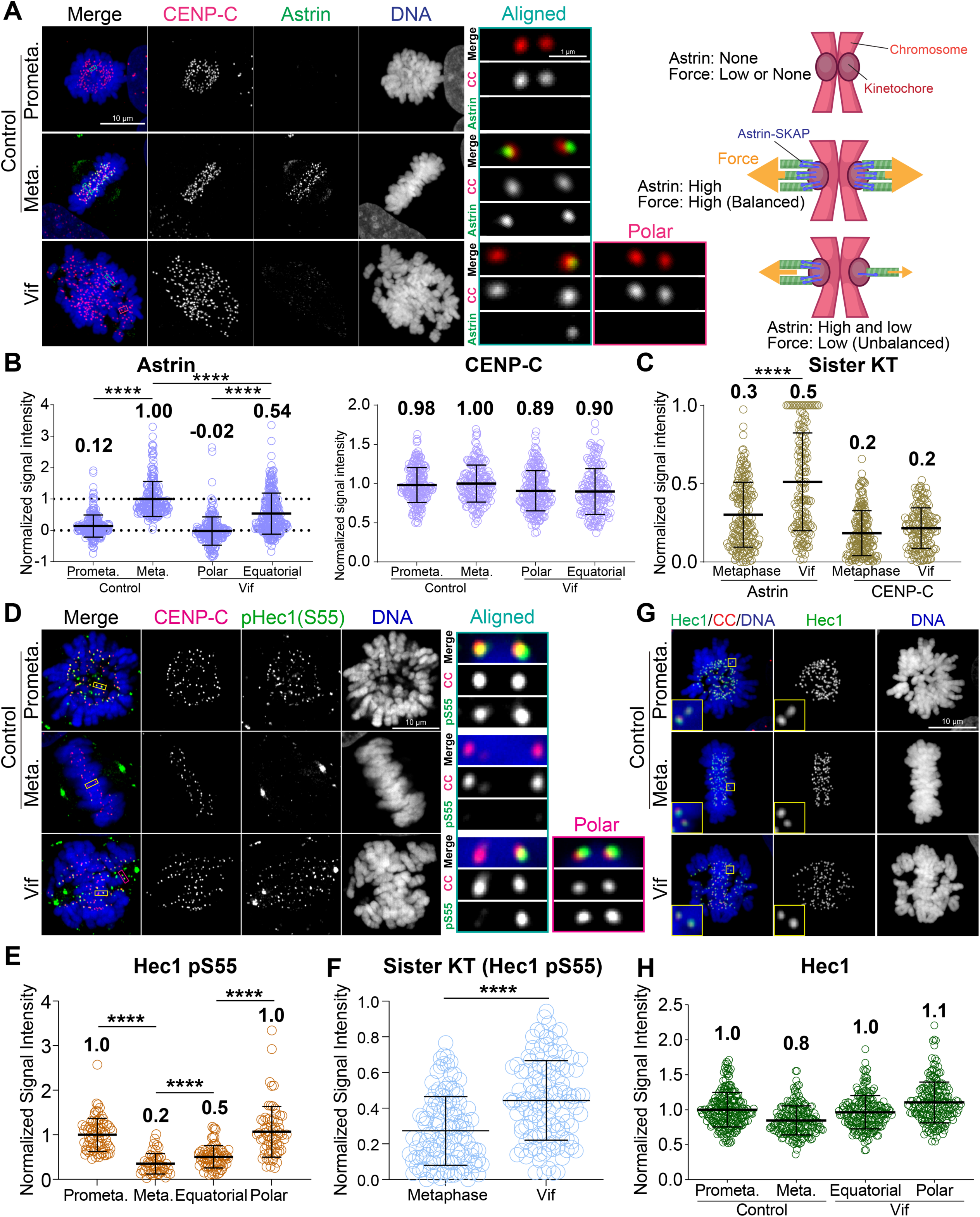
Vif impairs stable and balanced kinetochore microtubule attachments. (A) Left: Representative maximum intensity projection of fixed Control and Vif-expressing HeLa cells immunofluorescently labeled for CENP-C, Astrin, and DNA along with representative sister kinetochore pairs, Right: Illustrative interpretation of images on the left. (B) Average Astrin and CENP-C level at kinetochores for cells in (A) (n = 200 kinetochores from 8 cells from two independent replicates for each) (C) Comparison of Astrin and CENP-C on sister kinetochores, values normalized with formula: 1 – (lower intensity value/higher intensity value). (D) Representative image of fixed Control and Vif-expressing HeLa cells immunofluorescently labeled for CENP-C, pHec1(S55), and DNA along with representative sister kinetochore pairs. (E) Average pHec1(S55) level at kinetochores for cells in (D). (n = 200 kinetochores from 8 cells from two independent replicates for each). (F) Comparison of pHec1(S55) on sister kinetochores, values normalized with formula: 1 – (lower intensity value/higher intensity value). (G) Representative fixed-cell immunofluorescent image of HeLa cells expressing Vif, immunofluorescently labeled for Hec1, CENP-C, and DNA. (H) Average Hec1 level at kinetochores for cells in (G) (n=200 kinetochores over 8 cells from two independent replicates for each).

Generating uniform pulling force across sister kinetochores is essential for maintaining chromosome alignment at the cell equator during metaphase. While control cells showed equal Astrin recruitment at sister kinetochores, consistent with balanced forces (**Fig 6C)**, Vif-expressing cells showed significant differences in Astrin levels between sister kinetochores despite CENP-C levels remaining consistent (**Fig 6B-C)**.

The N-terminal domain of Hec1 has multiple phosphorylation sites, and dephosphorylation specifically by PP1 is critical for stabilizing its binding to microtubules (DeLuca *et al*, 2011). To directly validate the reduced activity of PP1 at kinetochores in Vif-expressing cells, we performed qIF using a phospho-Hec1 S55 antibody (pS55). As expected, phosphorylation levels pS55 were high in Control prometaphase and significantly reduced in metaphase (**Fig 6D-E**). In contrast, pS55 levels remained significantly high at aligned chromosomes (equatorial) in Vif-expressing cells compared to aligned metaphase chromosomes in Control cells (**Fig 6D-E**). Similarly, unaligned chromosomes (Polar) maintained pS55 levels similar to those in Control prometaphase. We confirmed that Hec1 levels at kinetochores were the same in both Vif-expressing and Control cells (**Fig 6G-H**). These results demonstrate that PP1 activity at kinetochores is weaker in Vif-expressing cells compared to Control cells. In agreement with the unbalanced Astrin recruitment at sister kinetochores in Vif-expressing cells, pS55 levels between sister kinetochores were also significantly unbalanced in Vif-expressing cells compared to Control cells (**Fig 6F**). In summary, Vif disrupts the proper assembly of the Astrin-PP1 complex at kinetochores, resulting in the retention of high phosphorylation Hec1 levels. This leads to weakened and uneven forces between sister kinetochores, likely contributing to dynamic chromosome movements.

### Limitations of the study

To study the spatiotemporal regulation of Vif and the effects of its expression levels at the single-cell level, we aimed to visualize Vif’s trafficking during the cell cycle. We discovered that C-terminal fusion of tags, such as 3xHA or mCherry, abolishes Vif’s ability to induce pseudo-metaphase arrest (**Fig S5A-E**). In this study, we elucidate the mechanisms underlying Vif-induced pseudo-metaphase arrest by utilizing cancer cell lines and a non-transformed normal cell line. While performing similar high-temporal resolution long-term imaging on well-established host cell types for HIV-1 (primary CD4+ T cells, lymphocytes, dendritic cells, or macrophages) poses significant technical challenges, future studies are required to investigate these cell types. Such investigations will help determine whether Vif can contribute to the suppression of the host immune system by effectively induce robust pseudo-metaphase arrest, ultimately leading to cell death.

## Discussion

The specific processes by which HIV-1 causes loss of CD4+ T cells are numerous and include activation of innate immune sensors (Doitsh & Greene, 2016), Envelope-driven cell fusion/syncytiation (Nardacci et al, 2015), and induction of cell cycle arrest followed by programmed cell death mediated by viral gene products that include Vif and Vpr (Muthumani et al, 2005). Vif has recently been shown to induce cell cycle arrest in conjunction with its downregulation of PP2A-B56 subunits (Marelli et al., 2020; Nagata et al., 2020; Salamango et al., 2019). However, the specific nature of this arrest was not previously examined at the level of single cells and had been assumed to occur during G2 based on flow cytometric assays. In our study, we discovered that expression of Vif actually reduces the duration of G2 (**Fig 3B**) and instead triggers a robust pseudo-metaphase arrest, confirmed in a broad range of cell lines, and with cells typically succumbing to apoptotic cell death after extended pseudo-metaphase (**Fig 1C, 1H, S1E, S2C**). We also demonstrate that, contrary to a prior study (Izumi et al., 2010), Vif-induced pseudo-metaphase arrest occurs independently of p53 status (**Fig 3D-F and S3D-F**).

Further, we demonstrate that Vif specifically disrupts the kinetochore functions, impairing proper mitotic progression (**Fig. 6A-B)**. Normally, after NEBD, microtubules efficiently capture kinetochores during prometaphase through the interplay of PP1 and PP2A-B56 phosphatase activities (Sivakumar & Gorbsky, 2017; Smith *et al*, 2019; Vallardi *et al*., 2017). PP2A-B56 is recruited to kinetochores in prometaphase, reducing Plk1 activity to facilitate kinetochore-microtubule assembly and promoting recruitment of PP1 by multiple adaptors. A major PP1 adaptor for kinetochore recruitment is the Astrin-SKAP complex whose recruitment requires proper microtubule end-on attachment (Conti *et al*., 2019; Friese *et al*, 2016; McVey *et al*, 2021). In Vif-expressing cells, Vif significantly reduces the level of PP2A-B56 at kinetochore in prometaphase, likely due to its role in B56 degradation. This reduction leads to a slower establishment of metaphase plate (**Fig 7**). The significant loss of PP2A-B56 at kinetochores impairs the feedback control necessary for stabilizing microtubule binding. As a result, there is a significantly lower and uneven recruitment of Astrin-SKAP-PP1 complex to kinetochores, causing uneven pulling forces between sister chromatids that result in some chromosomes being prematurely pulled towards spindle poles prior to satisfying the mitotic checkpoint (**Fig 7D, Step 1**). Upon approaching the spindle poles, Aurora A, another key kinase that regulates mitotic error correction, phosphorylates the MTBDs of Ndc80/Hec1 and destabilizes kinetochore-microtubule attachment (**Fig 7D, Step 2**) (Anna *et al*, 2015; Barr & Gergely, 2007; Chmátal *et al*, 2015; DeLuca, 2017; DeLuca *et al*., 2011; Kettenbach *et al*, 2011). This destabilization of the kinetochore-microtubule attachment could explain why polar chromosomes in Vif-expressing cells lose Astrin signals at kinetochores (**Fig 6A-B**). Polar chromosomes are then transported back to the equator by polar-ejection forces (**Fig 7D, Step 3**) (Poser et al, 2019; Wandke et al, 2012). The repetition of this cycle accounts for the observed abnormal dynamics of chromosome movements in Vif-expressing cells.

**Figure 7:**
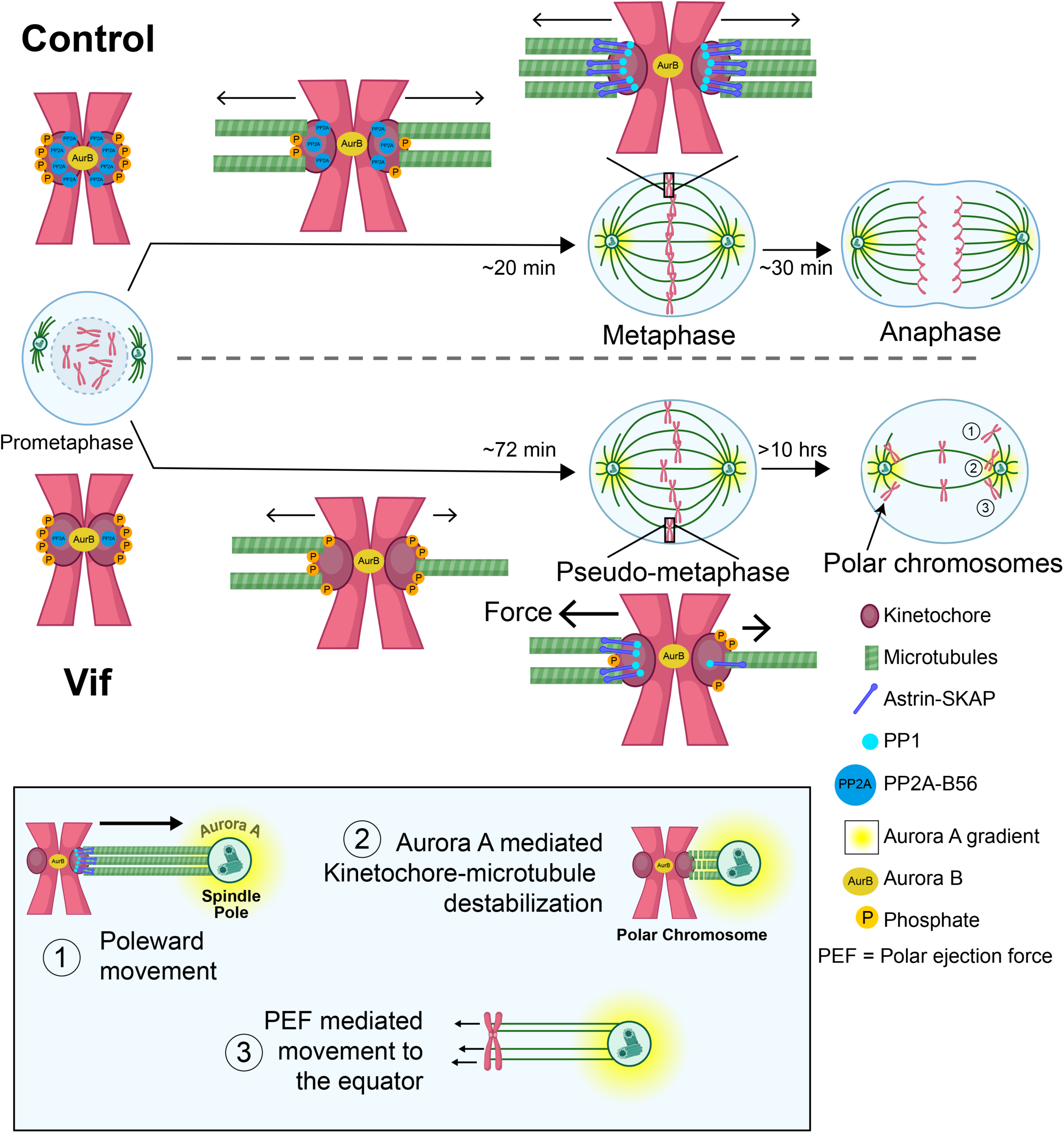
Proposed model for the molecular mechanism underlying Vif’s pseudo-metaphase arrest. Top: Cartoon model depicting metaphase alignment of Control cells followed by anaphase. Middle: Cartoon model depicting pseudo-metaphase alignment of Vif-expressing cells with unbalanced microtubule attachment followed by three-step polar chromosome cycle. Bottom: Cartoon depiction of three-step polar chromosome cycle, (1) chromosome at the equator is pulled towards pole due to unbalanced pulling force, (2) kinetochore-microtubule destabilization at the spindle pole, (3) equator-directed movement of chromosome for realignment.

## Acknowledgement

We would like to thank YuHi Hara, Takanori Tsuchiya, Yokogawa Corporation, Nikon, and Tokai Hit for critical equipment and technical support. We also would like to thank Dr. James Bruce for the critical suggestions and experimental support and Ms. Ainslie Homan for support in data analysis. Part of this work is supported by the University of Wisconsin-Madison Office of the Vice Chancellor for Research with funding from the Wisconsin Alumni Research Foundation, start-up funding from University of Wisconsin-Madison SMPH, UW Carbone Cancer Center, and McArdle Laboratory for Cancer Research, and NIH grant R35GM147525 and U54AI170660 (to A.S.) and U54AI170660, R01AI110221, and P01CA022443 (to N.S.).

## Author contribution

E.L.E. III initiated the project, identified Vif-induced metaphase arrest, and generated key viral reagents and NL4-3 strain HIV-1 reporter vectors. D.G. conducted the bulk of the follow-up precision imaging experiments and analyses, with assistance from M.B., K.G.S., N.S., and A.S.. A.S. and N.S. conceptualized, supervised and funded the project. D.G. and A.S. prepared the initial manuscript draft, with assistance from N.S., E.L.E. III, and M.B. All authors reviewed and contributed to the manuscript’s refinement.

## Methods

### Cell Culture

Human HeLa, RPE1, Cal51, and MDA-MB-231 cells were originally obtained from the American Type Culture Collection (ATCC, Manassas, VA, USA). RPE1 p53 KO, HCT116 p53 KO, RPE1 (H2B-RFP), MDA-MB-231 (H2B-mCherry), and Cal51 (Tubulin-mNeonGreen and H2B-mScarlet were endogenously tagged by CRISPR-Cas9) cells were originally obtained from Dr. Jan Korbel, Dr. Yue Xiong (UNC), Dr. Mark Burkard, and Dr. Beth Weaver, respectively. H2B-GFP expressing HeLa cells and conditional CO-Vif-expressing HeLa cells using a pCEP4 vector (Thermo) containing the TRE and Tet promotor with codon-optimized Vif or mNeonGreen were generated in this study. HeLa, MDA-MB-231, HCT116, RPE1 and Cal51 were grown in DMEM high glucose (Cytiva Hyclone; SH 30243.01) or DMEM/F12 (Cytiva Hyclone; SH 3026101) supplemented with 1% penicillin-streptomycin, 1% L-glutamine, and 10% fetal bovine serum under 5% CO_2_ at 37°C in an incubator.

### Live cell imaging

RPE1, Cal51, HeLa, and MDA-MB-231 cells were plated on 4-chamber 35mm glass bottom dishes at least one day prior to imaging (4 chamber with #1.5 glass, Cellvis). In a subset of experiments, cells were stained using sirDNA (Cytoskeleton, 150 nM) for 2 hours prior to imaging to visualize chromosomal DNA. For conditionally Vif-expressing cells, doxycycline (1 µg/ml, Sigma) was supplemented prior to imaging. High temporal resolution live-cell imaging was performed using a Nikon Ti2 inverted microscope equipped with a Hamamatsu Fusion camera, spectra-X LED light source (Lumencor), Shiraito PureBox (TokaiHit), and a Plan Apo 20x objective (NA = 0.75) controlled by Nikon Elements software. Cells were recorded at 37°C with 5% CO2 in a stage-top incubator using the feedback control function to accurately maintain temperature of growth medium (Tokai Hit, STX model). Images were recorded for 48-120 hours at 6-12 min intervals with three to four z-stack images acquired at steps of 1.5∼2 μm for each time point.

### Fixed high- and super-resolution imaging

HeLa cells infected “Control” (Vif-negative) and Vif-containing viruses were fixed by 4% PFA (Sigma) at 72 hours post-infection. Cells were then permeabilized by 0.5% NP40 (Sigma) and incubated with 0.5% BSA (Sigma). Following primary and secondary antibodies were used; CENP-C (MBL), Tubulin (Sigma), GFP (Thermo), B56-alpha (BD Biosciences), Plk1 (Santa Cruz), Astrin (Sigma), Hec1/9G3 (Abcam), Hec1 pS55 (GeneTex), anti-mouse IgG Alexa 488 (min X, JacksonImmuno research), anti-guinea pig IgG Rhodamine Red X (min X, Jackson immune research), anti-guinea pig IgG Alexa 647 (min X, JacksonImmuno research), anti-rabbit IgG Alexa-488 (JacksonImmuno research) and anti-rabbit Alexa 647 (min X, Jackson immune research). Stained samples were imaged both CSU W1 spinning disc confocal and CSU W1 SoRa super-resolution (Yokogawa) confocal microscope (Loi et al, 2023). These spinning disc confocal units were equipped with a Nikon Ti2 inverted microscope with a Hamamatsu Fusion camera, Shiraito PureBox (TokaiHit), and a TIRF SR 100x objective (NA = 1.49). The microscope system was controlled by Nikon Elements software (Nikon). Figure 3B images were generated using Imaris software (Andor).

### Image analysis

Image analysis was performed using Nikon Elements software (Nikon) or Metamorph (Molecular Devices). Mitotic stages and errors were determined by nuclear staining. The mitotic duration was defined as the time from nuclear envelope breakdown (NEBD) to anaphase onset. Timepoints of formation and loss of metaphase plate were documented. CFP signals were used as a marker for infected cells. Tubulin-mNeonGreen was used for quantifying numbers of spindle poles and monitoring their dynamics. Spindle pole distance was measured when spindle poles were maximally stretched in high-temporal live cell images using Nikon Elements.

### Cell cycle phase analysis

To track cell cycle progression, H2B signals were measured over time using Nikon NIS Elements on time lapse images of Cal51 cells. Signal intensities were measured manually and the local background correction method (Loi *et al*., 2023; Suzuki *et al*, 2015) was applied to accurately quantify chromatin signal intensity. Signals were collected in this manner at 18-30 min intervals. The duration of each cell cycle stage was determined by analyzing changes in the H2B-mScarlet signal over time.

### Polar chromosome quantification

Cell segmentation and measurements of chromosome distribution were performed using the Nikon NIS Elements program. First, the region-of-interest (ROI) tool was used to select chromosomes located at each pole or at the equator. Corrected signal intensity was calculated using a local background correction method (Suzuki *et al*., 2015). Measurements were made for every 6 minutes for the first 2 hours after NEBD, and for every ∼1.5 hours subsequently. For each time point, the percentage of polar chromosomes was calculated using the following formula: (Corrected intensity of Pole1+Pole2)*100/(Corrected intensity of Pole1+Pole2+Equator).

### Spindle rotation measurements

Measurements of spindle rotation were performed using the Nikon Elements program’s Manual Measurement tool. Cells were observed after NEBD and the free angle tool was used to measure the absolute value of the spindle rotation angle traced from using either the spindle pole or the equatorial chromosomes as a reference. For control cells, measurements were made for each consecutive frame from the first frame where a spindle appeared until the first frame at the onset of anaphase. For Vif-expressing cells measurements were made for frames whenever a visually significant angle was traced. Data was exported to Excel. The Matplotlib library in Python was used to make polar plots, with time plotted as radius and angle traced plotted as theta.

### Statistics

All experiments were independently repeated 2-3 times for mitotic duration measurements. p-values were calculated using one-way ANOVA and the two-tailed Student’s t-test. p-values < 0.05 were considered significant. In the figures, p-values are denoted as * for ≤0.05, ** for ≤0.01, *** for ≤0.001 and **** for ≤0.0001.

### Transduction and infection

For infections, growth media was replaced with viral supernatants carrying VSV-G-pseudo typed HIV-1 CFP reporter viruses (Vif-positive or Vif-negative) at a multiplicity of infection of ∼1, with the viruses engineered and produced as previously described (Evans et al., 2018).

